# JASMONIC ACID PRIMING OF POTATO USES HYPERSENSITIVE RESPONSE-DEPENDENT DEFENSE AND DELAYS NECROTROPHIC PHASE CHANGE AGAINST *PHYTOPHTHORA INFESTANS*

**DOI:** 10.1101/2021.03.19.436215

**Authors:** Diego F Arévalo-Marín, Daniel M Briceño-Robles, Teresa Mosquera, Luz Marina Melgarejo, Felipe Sarmiento

## Abstract

One of the most important diseases affecting potato is late blight, caused by the oomycete *Phtytophthora infestans*. The use of jasmonic acid has been reported to reduce the progression of the disease in potato, but the defense mechanisms involved in this response are unknown. In this study we described the effect of jasmonic acid as a priming agent over time in the defense response of potato against the invasion of *P. infestans*. We observed that the initial stimulus generated by the exogenous application of jasmonic acid had an effect on the stomatal conductance of the treated tissue and activated *StMYC2* expression. Results reveal a priming effect in plants inoculated 11 days after treatment with jasmonic acid, evidenced by an increased transcriptional induction of defense-associated genes, decrease in the number of necrotic lesions and an evident reduction of lesion area (72.23%). Furthermore, in this study, we show that the tested concentration of jasmonic acid does not have an adverse effect at the physiological level in plants, since variation in stomatal conductance was transient, no change in chlorophyll a fluorescence and no early senescence in leaves was observed.

## 1. Introduction

Potato (*Solanum tuberosum*) is the third most consumed food crop in the world [1]. This species is affected by the pathogen *Phytophthora infestans* (Mont.) de Bary causing late blight disease, considered the main phytosanitary problem for potato production [2,3]. This disease can generate yield losses of around 40%, even up to 100% in field conditions in susceptible varieties. These losses represent an annual financial loss of approximately € 6 billion [4,5].

*Phytophthora infestans (P. infestans)* is a hemibiotrophic oomycete with a two-phases infection style. An initial biotrophic infection phase, where the pathogen requires living cells, which is followed by a second necrotrophic phase. In the necrotrophic phase, the pathogen feeds on dead plant tissue. The change of phase occurs at around 24 – 48 hours post infection [6]. The response of the plant against *P. infestans* causes notable changes at the physiological level, such as reduced photosynthesis, changes in transpiration, changes in membrane permeability, increased respiratory rate, and changes in tissue expression profiles among others [7,8,9,10].

The response of plants to biotic stresses is influenced by signaling pathways regulated by plant hormones such as jasmonic acid (JA) and salicylic acid (SA). These hormones coordinate defense induction in plants depending on the type of pathogen [11]. JA activation is generally associated with defense responses against necrotrophic pathogens, as well as with establishment of systemic induced resistance (ISR); on the contrary, biotrophic and hemibiotrophic pathogens responses are dependent on accumulation of SA and are associated with establishment of systemic acquired resistance (SAR) [12,13,14]. However, evidence accumulate towards the involvement of jasmonic acid in the response to biotrophic pathogens [15,16] and hemibiotrophic pathogens [17,18,19].

Plant resistance against pathogens may be induced by exposure to an exogenously applied chemical stimulus. The application of chemical compounds acts as a priming stimulus preparing signaling pathways downstream to respond fast and strong, thus improving the defense response of plants against future biotic stresses [20,21,22]. Various naturally occurring chemical compounds such as: ethylene (ET), salicylic acid (SA), jasmonic acid (JA) and abscisic acid (ABA) and some non-protein amino acids such as: β-aminobutyric acid (BABA) and pipecolic acid have been reported as defense priming agents [23].

Priming and defense responses differ between plant species and the priming stimulus. These compounds enhance the transcriptional activation of defense-associated genes, the accumulation of biologically active signals or molecules (mRNA, amino acids, phenylpropanoids), improvement of the cell wall structure, generation of reactive oxygen species (ROS), among others [21,24,25,26]. Previous studies have shown that application of JA act as a priming stimulus by improving the defense response in various species, incrementing the expression of defense genes against attacks on hemibiotrophic pathogens [17,18,27,28].

Research towards priming for potato response to biotic and abiotic stress report effects of various molecules [29,30,31]. However, the short and medium-term side effects have not been thoroughfully analyzed. Several of these compounds interact with primary metabolism, and can affect plant growth and development [31,32]. In order to detail the effect of JA in the plant and further describe its role as a priming agent in potato cv. Criolla Colombia against *P. infestans*, we initially monitored the plant response to JA during 17 days. Then we tested the time required for the plant to achieve priming by inoculating at three different moments during the duration of the experiment, and analyzed the plant responses during challenge with the pathogen at a phenotypical, physiological and transcriptional level. Our results show a priming effect in plants inoculated 11 days after treatment with jasmonic acid, evidenced by a reduction in lesion area, decrease in the number of necrotic lesions and an enhanced transcriptional induction of defense-associated genes.

## 2 Materials and methods

### 2.1 Plant material and growing conditions

Certified tubers of the diploid commercial variety “Criolla Colombia” (*Solanum tuberosum* Group *Phureja*), susceptible to *P. infestans* were planted in pots of 20 cm in diameter. A mixture of soil and sand in a ratio of 3:1 was used [33]. The plants grew under greenhouse conditions in an air temperature range between 17 to 20 °C with light: dark cycle of 12:12h and a general average air relative humidity of 55%. During the course of the experiment, the plants were watered three times per week with running water at field capacity. Plants were kept in optimal conditions of nutrition and sanitary status.

### 2.2 Jasmonic Acid treatment

When the plants had 10 fully extended leaves from the main stem (stage 110 BBCH scale) [34], the three terminal leaflets of three leaves from the middle third of ten plants were sprayed on the adaxial surface with 900 uL of a solution of JA (Jasmonic Acid, Liquid Bioreactive SIGMA -JA), each leaflet was sprayed with 100 uL at a concentration of 150 μg mL^-1^ plus tween 80 at 0.01% using an atomizer [31]. The solutions were prepared from a 100 mg JA stock solution which was dissolved in 1000 uL of 96% ethanol (w/v). A plastic bag was used at the time of the application to avoid drift to other leaves and plants.

### 2.3 Culture and inoculation with Phytophthora infestans

An isolate of *P. infestans* (clonal lineage EC-1, mating type A1), collected by Prof. Mosquera’s lab in the municipality of Sibate, Colombia (An important area for potato production), was used. *P. infestans* cultures were grown on rye agar at 18 °C in dark conditions. The isolate was periodically activated by placing infected plant tissue on thin slices of potato tubers of a susceptible variety (Tuquerreña) to maintain virulence and abundant and fresh sporulation. The mycelium was constantly transferred to rye agar culture media to maintain purity [35,36,37]. The inoculum for the assays was prepared by scraping the sporangia from 15-20 day-old rye agar media with sterile water. The sporangia concentration was determined using a Neubauer camera and adjusted to a concentration of 50,000 sporangia mL^-1^ [38]. For inoculation, a 10 μL drop of a suspension of *P. infestans* sporangia was inoculated on the abaxial side of the terminal leaflets using a HandyStep™ S micropipette (BRAND™ 705110). After inoculation, the plants were covered with transparent plastic bags sprayed with sterile water inside to maintain high humidity and facilitate infection. Plastic bags were removed 48 hpi.

### 2.4 Experimental design

A factorial design with two factors and three levels per factor was used. The two factors consisted of JA application and application of mock treatment (MT) without JA (H_2_0 + tween 80 at 0.01% + ethanol at 20%). Plants with 10 unfolded main stem leaves were treated with of JA by spraying at a concentration of 150 μg mL^-1^. The three levels consisted of inoculation of *P. infestans* 5, 11 and 17 days after treatment (DAT). In parallel, plants were treated with MT and inoculated with *P. infestans* 5, 11 and 17 DAT. A summary of the experimental design used is shown in Figure S1A. Ten experimental units were used for each treatment, each consisting of three terminal leaflets of three leaves from the middle third.

To control the effect of JA on potato plants, an independent experiment was set in which plants were only treated with JA or MT. Plants were placed in a completely randomized design. A summary of the experimental design used is shown in Figure S1B. Each experimental unit consisted of three terminal leaves of three leaves from the middle third; ten experimental units were used for each treatment.

### 2.5 Measurement of chlorophyll a fluorescence

The fluorescence of chlorophyll was estimated using a portable non-modulated pulse fluorometer (Pocket PEA, Hansatech Instruments, UK). Measurements of *P. infestans* inoculated treatments were taken every 48 hpi for a period of 8 days, thus, at 48, 96, 144 and 192 hpi. Measurements for uninoculated treatments were taken every two days for a period of 16 days, thus, 2, 4, 6, 8, 12, 14 and 16 DAT. Measurements of the treatments were made at pre-dawn (04:00 – 05:00) on 10 plants in the leaflets of three leaves per plant from the middle third. The initial fluorescence (F_0_), and maximum fluorescence emission (F_m_) parameters were measured to calculate the maximum quantum efficiency of photosystem II photochemistry according to the equation (1) [39].

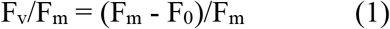

### 2.6 Measurement of stomatal conductance and leaf temperature

Variables related to stomatal conductance (g_s_; mmol H_2_O/m*s^2^) and leaf temperature (°C) were estimated using a portable leaf porometer (SC-1, Decagon Services Inc, WA, EE. UU). Measurements of treatments inoculated with *P. infestans* were made every 48 hpi for a period of 8 days, thus, at 48, 96, 144 and 192 hpi. Measurements for uninoculated treatments were taken every two days during a period of 16 days, thus, 2, 4, 6, 8, 12, 14 and 16 DAT on 10 plants on the same leaflets used to measure of chlorophyll a fluorescence. Measurements were made between 08:00 and 10:00 h.

### 2.7 Disease progression assessment

Disease progression assessment was carried out by measuring the area of the lesion with a digital caliper (Truper). Measurements were made every 48 hpi over a period of 8 days, thus, at 48, 96, 144 and 192 hpi. The Area Under the Disease Progress Curve (AUDPC) was calculated according to the procedure of the International Potato Center, using the equation proposed by Campbell and Madden (1990). Relative AUDPC was calculated according to equation (2) to compare between treatments [40,41].

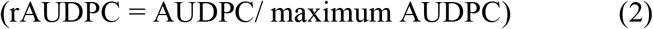

### 2.8 Analysis of the infection process by light microscopy -Trypan blue staining

The infection of *P. infestans* in plants treated with JA or MT was performed as described previously. Samples of infected leaves were collected every 24 hpi during 3 days, at 24, 48 and 72 hpi, using a cork borer (diameter 7mm). Four discs were collected per evaluation time for a total of 12 discs per treatment. The disks were rinsed in an acetic acid-ethanol clearing solution, (1:3 *v*/*v*) for 16 h, transferred to an acetic acid-ethanol-glycerol solution, (1:5:1 *v*/*v*/*v*) for at least 3 h and taken to a solution of trypan blue 0.01% (*w*/*v*) in lactophenol for 12 h. The discs were subsequently washed with 60% sterile glycerol and fixed to microscopy glass slides [42,43,44]. Images were taken using a light microscope (CX-31 Olympus Corporation Tokyo, Japan) and a digital camera. The images were analyzed using the software ToupView V.3.7. The number of necrotic lesions for each treatment were quantified by counting lesions in at least 3 visual fields per biological replicate.

### 2.9 Quantitative RT-PCR analysis

Samples treated with JA and MT, without inoculum, were taken only 2h, 5, 11 and 17 DAT, while for the inoculation treatments tissue was taken 48 hpi, the total RNA was extracted using the InviTrap® extraction kit. Spin Plant RNA Mini Kit (INVITEK) following the manufacturer’s protocol. Starting from 1 µg of purified total RNA, genomic DNA removal was performed using DNase I treatment (Thermo Scientific). First strand synthesis was accomplished using the RevertAid H Minus First Strand cDNA Synthesis Kit (Thermo Scientific) following the manufacturer’s protocol. Quantitative RT-PCR was performed using the gene-specific primers related to the jasmonic acid-mediated response (*StMYC2-Like – StMYC2; StChalcone synthase -StCHS*), with qualitative defense response (*StPR-1*) and genes previously associated with defense in potato (*StPeroxidase 2-like; StLipoxigenase -StLOX, StOsmotine, StWRKY1*) (Table S1) [45,46,47,48,49,50,51]. Data were normalized using the reference gene *Elongation Factor 1α* (*StEF-1α*) as it has been reported to be one of the most stable reference genes for potato data normalization during infection with *P. infestans* [45,52]. QRT-PCR reactions were carried out with SsoFastTM EvaGreen® Supermix with Low ROX (Biorad, USA) in the LightCycler® 96 kit (Roche) using 3 biological replicates for each of the 8 treatments, with triplicate technical replicates for each biological replicate. The relative quantification of the specific levels of mRNA in relation to the normalizing gene was calculated according to the method proposed by Muller et al. [53] using the Q-Gene software V.1.0 [54].

### 2.10 Statistical analysis

Physiological variable data were analyzed with MannWhitney-U and Kruskal-Wallis non-parametric test (P≤0.05 and P≤0.01) to detect differences among treatments. Relative gene expression data, rAUDPC and microscopy were subjected to the t-Student test at each time point to analyze differences between plants treated with JA and MT. The SPSS Statistics V.25.0 software program (SPSS Software Inc., Chicago, IL, EE. UU, 2016) was used for statistical analysis and the Graphpad Prism V.6.0 software to construct the graphs (GraphPad Software Inc., San Diego, CA, EE. UU, 2015).

## 3. Results

### 3.1 Jasmonic acid treatment modifies the physiological response of potato plants by reducing stomatal conductance

To demonstrate if JA modifies the physiological state of the plants, chlorophyll a fluorescence and stomatal conductance were evaluated for 16 days in plants treated with AJ and MT. The Fv/Fm (Figure 1A) presented a constant trend over time and no significant differences were observed between treatments at any of the times evaluated. The values of the maximum quantum efficiency of PSII photochemistry (Fv/Fm), during experiment were close to 0.836 Fv/Fm.

**Figure 1.**
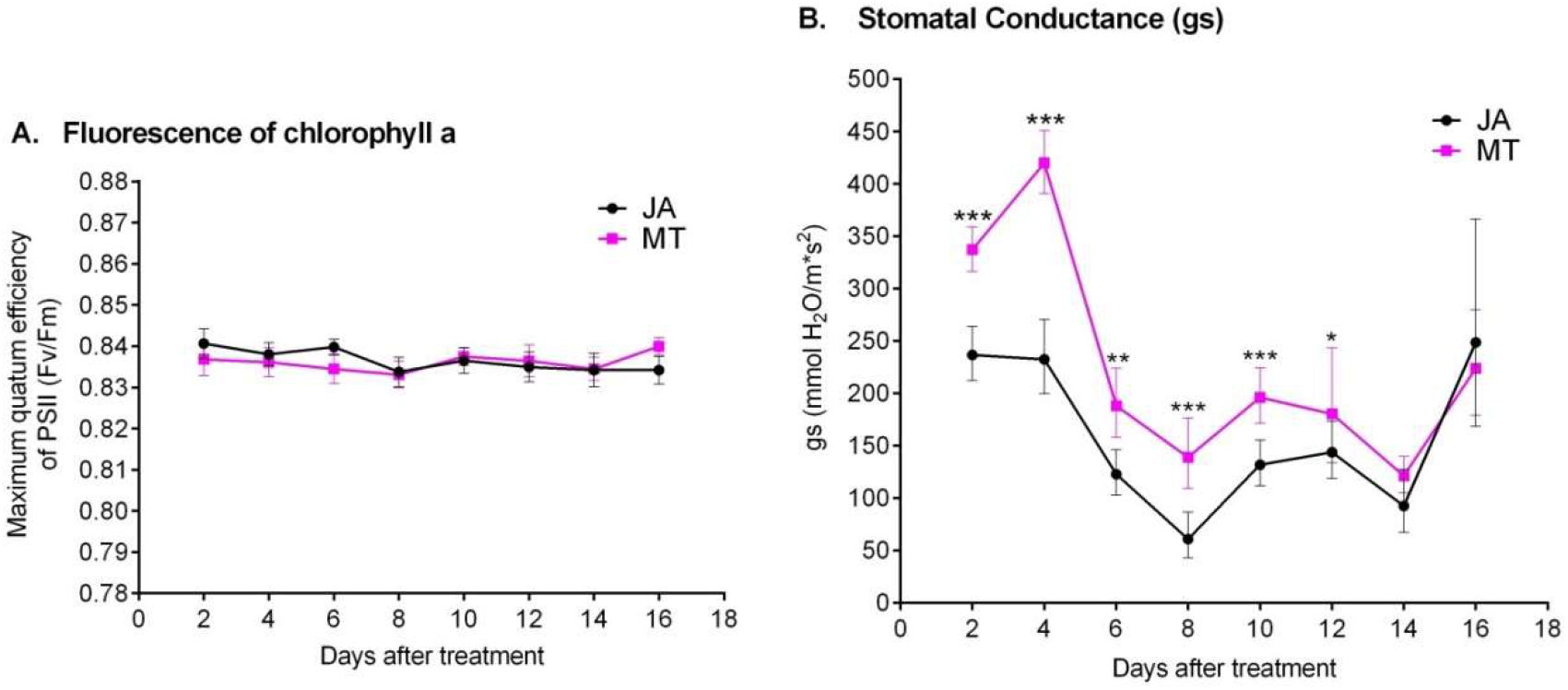
Physiological response of potato plants treated with jasmonic acid (JA) or mock treatment (MT). (A) Maximum quantum efficiency of PSII photochemistry (F_v_/F_m_) and (B) Stomatal conductance (g_s_). Measurements of physiological variables were taken every two days for 16 days after the application of jasmonic acid or mock treatment. Each point represents the average 10 plants ± SE. Values represent means of at least three independent experiments. Significant differences were calculated using the MannWhitney-U test. *: Significant at level <0.05, **: Significant at level <0.01, ***: Significant at level <0.001.

Stomatal conductance exhibited significantly lower values (P <0.01) in plants treated with JA compared to plants treated with MT until 12 DAT. At 10 DAT (p<0.001^***^), MT-treated plants displayed a mean stomatal conductance of 196,273 mmol H_2_O/m*s^2^ compared to JA-treated plants with a mean of 131,882 mmol H_2_O/m*s^2^, which corresponds to a 32.81% reduction (Figure 1B). After 14 DAT (p=0.364), no significant differences were observed between the two treatments.

### 3.2 Exogenous application of jasmonic acid protects potato leaves by slowing down the growth of P. infestans

The effect of JA on the progress of *P. infestans* infection was evaluated by means of rAUDPC analysis among treatments. Figure 2 reveals a significant decrease (72.23%; p<0.001***) of rAUDPC in inoculated JA-treated plants 11 DAT compared with MT-treated plants. The reduction of rAUDPC in inoculated JA-treated plants 5 DAT was 33.18% and a greater variability was observed in plants treated with the hormone (p=0.277). At 17 DAT, no significant difference (p=0.847) in lesion progression was observed between the two sets of plants. These findings suggest the ability of JA to interfere with the progress of *P. infestans* in potato.

**Figure 2.**
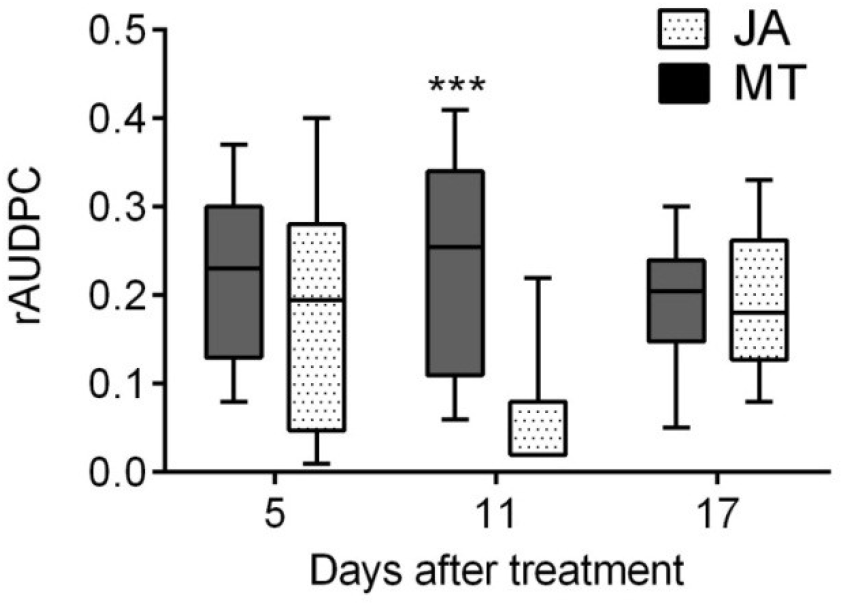
Effect of jasmonic acid on the progress of the disease. rAUDPC measurement was calculated from percentage of diseased area taken every 48 hours for eight days after inoculation with *P. infestans* on potato plants previously treated with jasmonic acid or mock treatment. The data presented are the means ± SE of 10 plants by experimental condition. Values represent means from at least three independent experiments. Significant differences were estimated using the t-Student test, ***: Significant at level <0,001.

### 3.3 The physiological response of the plants to stress caused by P. infestans is modified by jasmonic acid pretreatment after 11 DAT

After the successful establishment of *P. infestans* in plants, the physiological response of the plants was evaluated at different time points after inoculation. Both stomatal conductance (g_s_) and the maximum quantum efficiency of PSII photochemistry decreased as the development of the disease progressed (Figure 3). The Fv/Fm exhibited significant differences (P <0.05) among treatments at 48 and 96 hpi, being JA-treated plants 11 DAT the one that presented significantly higher values with means of 0.831 (p=0.035*) and 0.711 (p=0.012*), respectively (Figure 3A). Stomatal conductance (Figure 3B) presented significant differences (P <0.05) at 144 and 192 hpi, in JA-treated plants 11 DAT, showing significantly higher values with means of 140.95 (p=0.005**) and 110.31 (p=0.002**) mmol H_2_O/m*s^2^, respectively. These results show that JA reduces the physiological stress of invasion by *P. infestans* particularly 11 days after its application.

**Figure 3.**
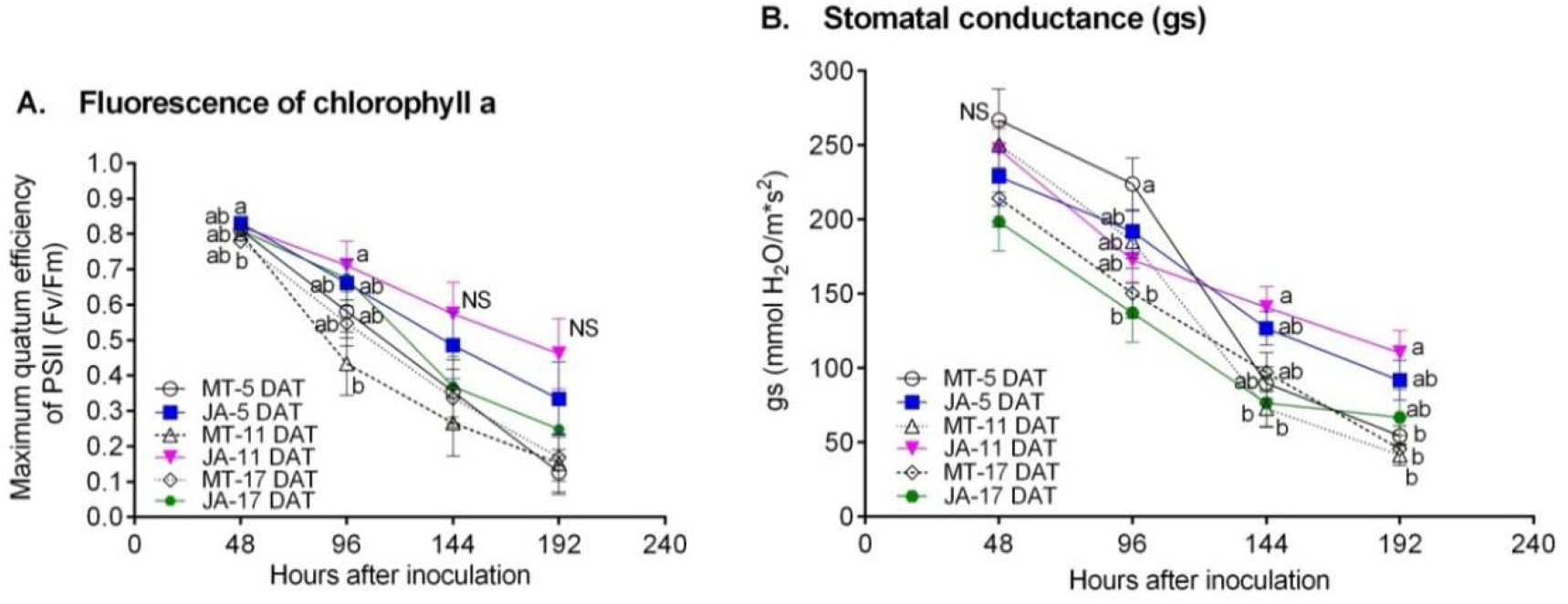
Physiological response of potato plants treated with jasmonic acid (JA) or mock solution (MT) and inoculated with *Phytophthora infestans*. **(A)** Maximum quantum efficiency of PSII phochemistry (F_v_/F_m_) and **(B)** Stomatal conductance (g_s_). Measurements of physiological variables were made every 48 hours for eight days after inoculation with *P. infestans* on potato plants previously treated with jasmonic acid or distilled water (control). Each point represents the average 10 plants ± SE. Values represent means from at least three independent experiments. Different letters correspond to significant differences between treatments (Kruskal – Wallis, p<0.05), NS: Not Significant.

### 3.4 Exogenous application of jasmonic acid delays the infection process of P. infestans and reduces the number of necrotic lesions

To reveal the differences in the infection cycle of *P. infestans* in plants after treatment with JA, a microscopy analysis was performed to compare the structures developed by the pathogen in plants inoculated 11 days after the application of JA and MT. A, lower zoospore release (Figure 4A, E), lower abundance of invasive hyphae (Figure 4B, F) and branched mycelium (Figure 4C, G) was observed in plants treated with JA compared with the MT treatment. At 72 hpi, dark brown necrotic zones were detected in the MT treatment (Figure 4D) while light brown zones were noticed in plants treated with JA, (Figure 4H). According to the data, the treatment with JA reduces the development of the disease, evidenced in a lower abundance of structures developed by the pathogen compared to plants treated with MT.

**Figure 4.**
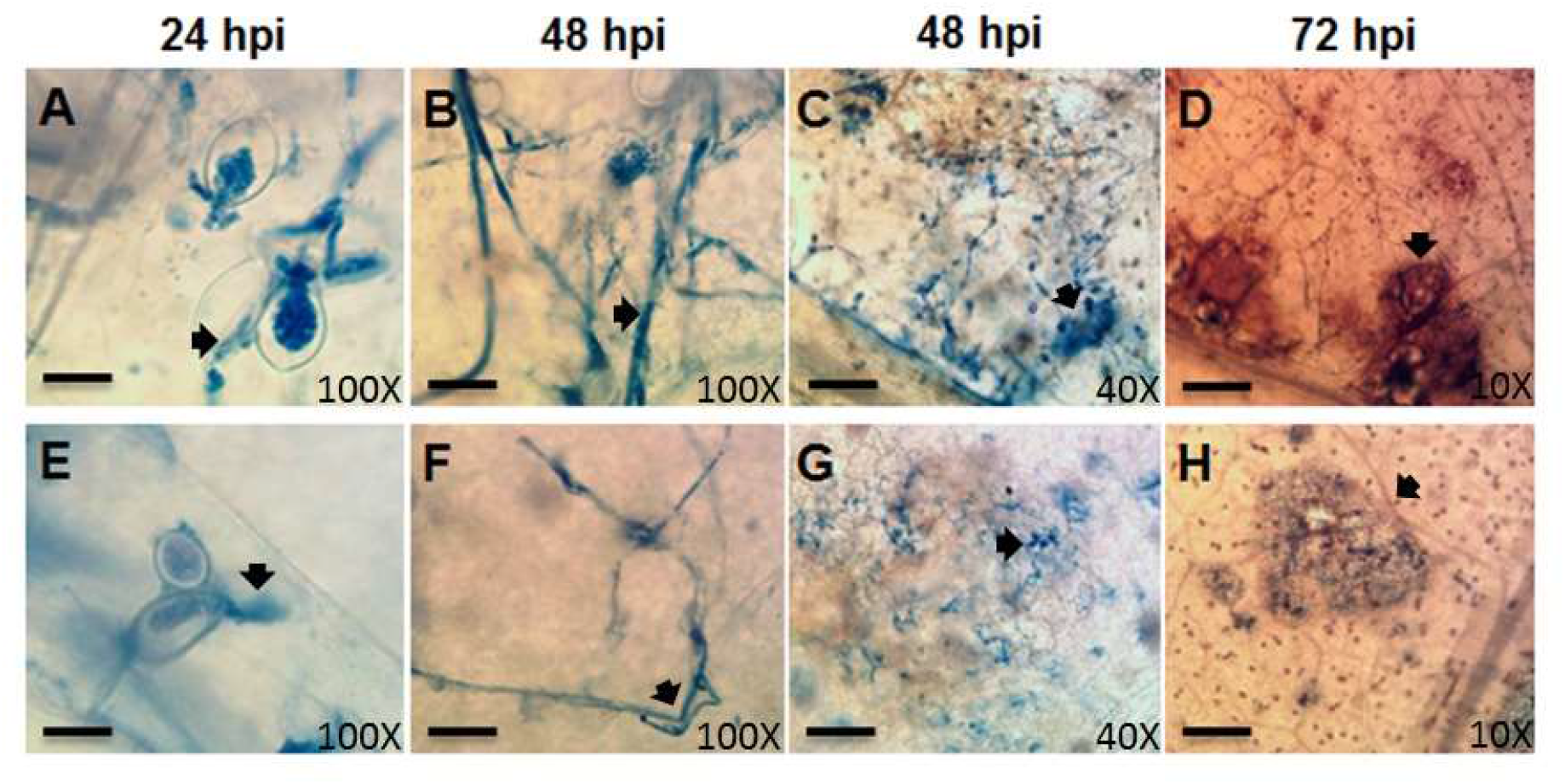
Microscopic assessment in potato treated with jasmonic acid (JA) or mock solution (MT) and inoculated with *P. infestans*. Development stages of the pathogen 11 days after the application of mock solution **(A-D)** and jasmonic acid **(E-H). (A, E)** Broken sporangia liberating zoospores (sp) were observed 24 hpi. **(B, F)** 48 hpi invasive hyphae (Hy) develops in both treatments. (**C, G**) Ramified mycelium (Rm) is observed 48 hpi. (**D, H**) Necrotic zones (Nz) are visible 72 hpi. Scale bars: (A, E, B, F) 20μm; (C, G) 50μm; (D, H) 100μm.

A significant reduction in the amount of necrotic lesions was observed in plants treated with JA and inoculated with *P. infestans* 5 and 11 DAT. At 5 DAT the JA treatment presented a mean of 6 lesions after 72 hpi, compared to 7.6 lesions of the MT treatment (P=0,042) (Figure 5A). The lowest amount of necrotic lesions was observed 11 DAT the JA treatment presented a mean values of 1.6 (at 48 hpi) and four lesions (at 72 hpi), compared to 4.5 (at 48 hpi; p=0.002**) and 6.8 (at 72 hpi; p=0.026*) lesions (Figure 5B) of the MT treatment. At 17 DAT, no significant differences were observed in the number of necrotic lesions between treatments (Figure 5C). In conclusion, JA reduces the progression of the disease by delaying the onset of the necrotrophic phase, especially in plants inoculated 11 DAT.

**Figure 5.**
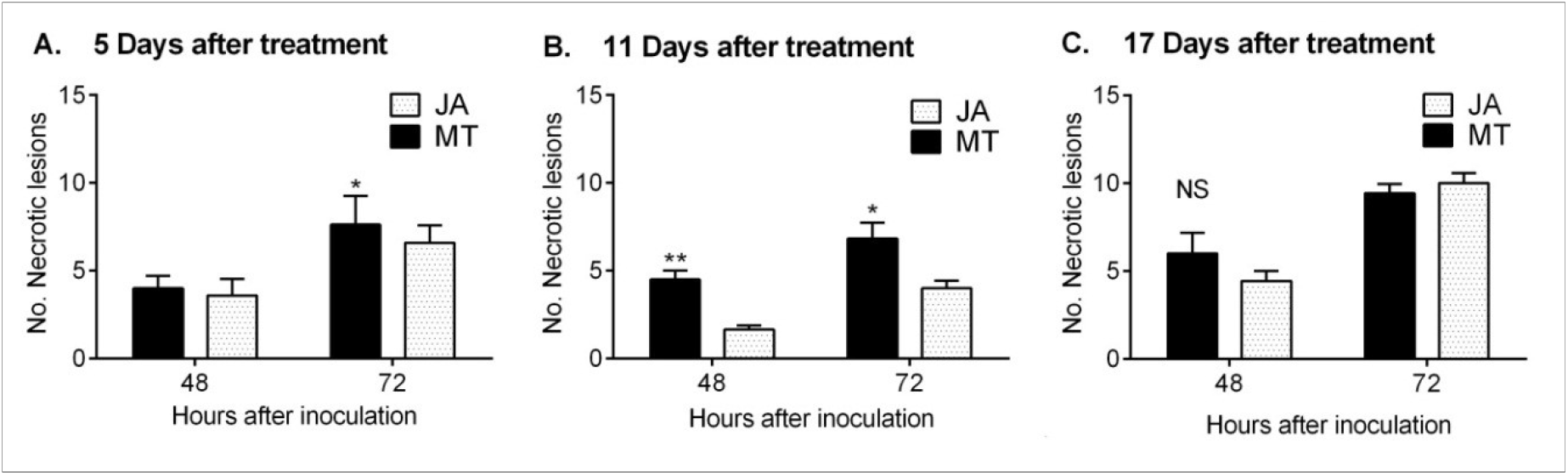
Necrotic lesion count in leaves infected with *P. infestans* 48 and 72 h after inoculation. **(A)** 5 DDT with JA or MT **(B)** 11 DDT with JA or MT, and **(C)** 17 DDT with JA or MT. The bars represent the mean ± SE. Values represent means from at least three independent experiments. Significant differences were estimated using the t-Student, NS: Not Significant, *: Significant at level <0.05, **: Significant at level <0.01.

### 3.5 StMYC2, a gene associated with jasmonic acid signaling is upregulated upon jasmonic acid treatment

To understand the effect of JA and to trace the downstream signaling process, the expression of *StMYC2*, a master transcription factor in the activation of signaling by JA was analyzed [55] at different time points after JA and MT application in plants not inoculated with *P. infestans*. Relative differences in transcriptional expression among treatments confirmed a significant increase in gene expression in plants treated with JA compared to MT (Figure 6). *StMYC2* displayed significantly higher relative expression levels (P <0.05) in JA-treated plants 2 hours after treatment (p=0.049*), 2 DAT (p=0.039*) and 5 DAT (p=0.033*), with levels being 3.58, 6.35 and 5.11 times higher compared to MT treatments. Transcriptional activation decreases at 11 days, where no significant differences were observed between application of JA and MT (p=0.906). These results support the idea that exogenous application of JA activates the JA-dependent signalling in local tissue, an effect that in the case of *StMYC2*, was observed to increase after two hours until 5 DAT.

**Figure 6.**
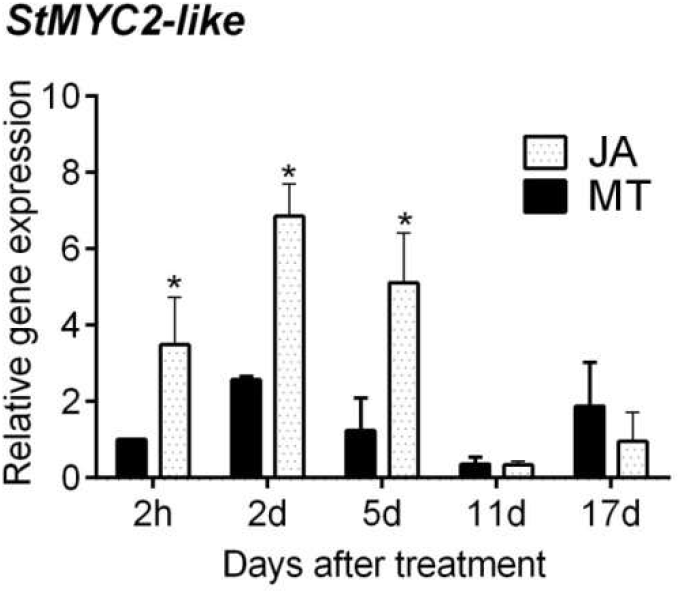
Relative transcriptional expression of *StMYC2-Like* after JA or MT treatment in potato plants. The relative expression was normalized with housekeeping *StEF1*α. Expression levels were calculated using the 2h MT treatment as a reference. The bars represent the mean ± SE. Values represent means from at least three independent experiments. Significant differences in relative expression levels using the t-Student, *: Significant at level <0.05.

### 3.6 Exogenous application of jasmonic acid increases the expression of genes associated with the jasmonic acid-mediated response in plants inoculated with P. infestans

To understand the priming effect of JA and inquire about the signaling network that is responsible for the protective effect against *P. infestans*, we analyzed the expression of genes associated with JA signaling. Samples were taken 48 hpi from inoculated plants treated with MT and JA 5, 11 and 17 DAT. Relative differences in transcriptional expression among treatments (Figure 7) confirmed, overall, an increase in gene expression in plants treated with jasmonic acid and inoculated with *P. infestans. StMYC2* (Figure 7A) exhibited a significantly higher relative expression (p=0.007**) in JA-treated plants inoculated 5 DAT, the level of expression being ten times higher compared to the MT treatment. The transcript of *StCHS* (Figure 7C) did not show significant differential expression among treatments. These results demonstrate that inoculation of *P. infestans* does not interfere with progression of the JA-associated signaling pathway.

**Figure 7.**
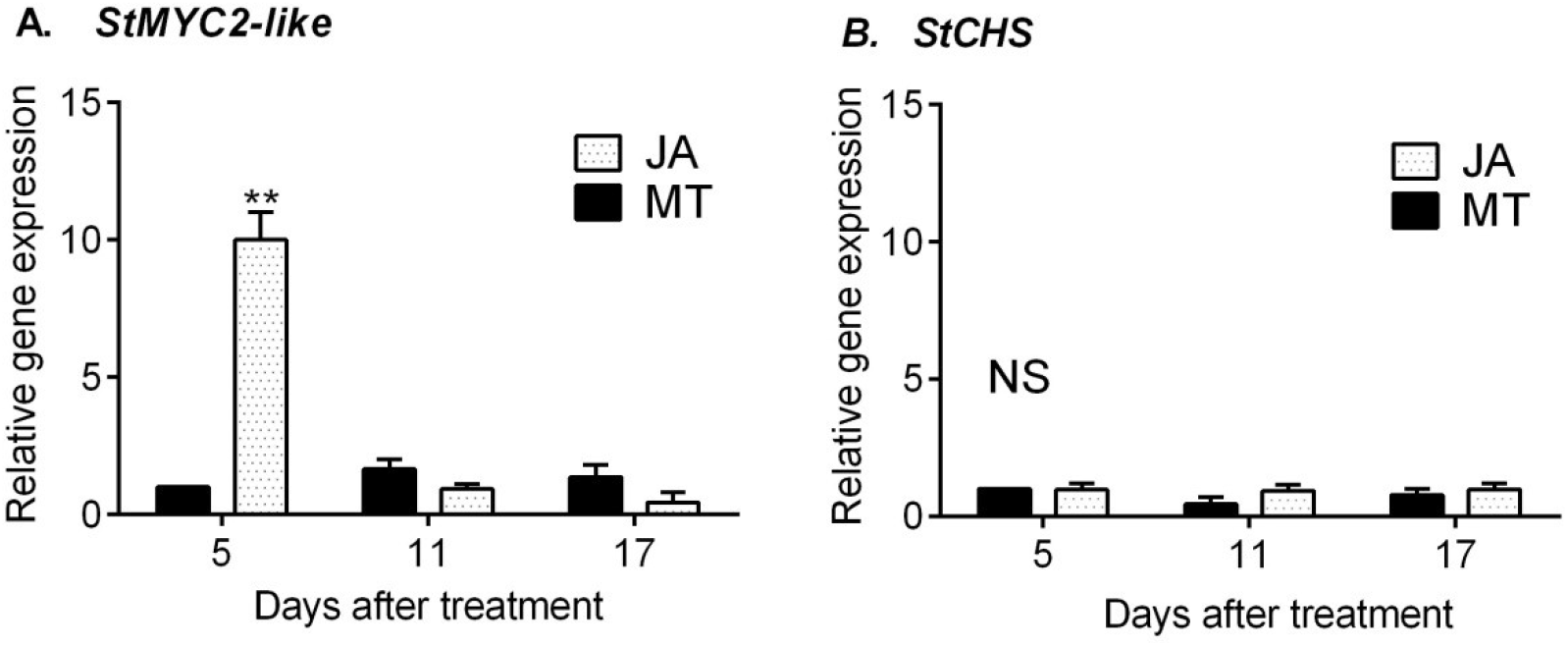
Relative transcriptional expression of JA-signaling response genes in potato treated with JA or MT and inoculated with *P. infestans* 5, 11 and 17 days after treatment (DAT). **(A)** *StMYC2-Like*, **(B)** *StCHS*. Samples were taken 48 hours post-inoculation. Expression was normalized with housekeeping *StEF1*α and relative expression levels were calculated using MT-5DAT treatment as a reference. The bars represent the mean ± SE. Values represent means from at least three independent experiments. Significant differences in relative expression levels using the t-Student, NS: Not Significant, *: Significant at level <0.05, **: Significant at level <0.01.

### 3.7 Exogenous application of jasmonic acid increases the expression of genes related to defense

Relative differences in transcriptional expression between JA and MT treatments generally exhibited higher gene expression in plants treated with JA and inoculated with *P. infestans. StPR-1* (Figure 8A) showed a significantly higher relative expression (P <0.05) in JA-treated plants inoculated 5DAT (0.023*) and 11 DAT (0.037*). Increments of three and five times were observed, compared to MT. At 17 days, no significant differences were found between plants treated with JA or MT.

**Figure 8.**
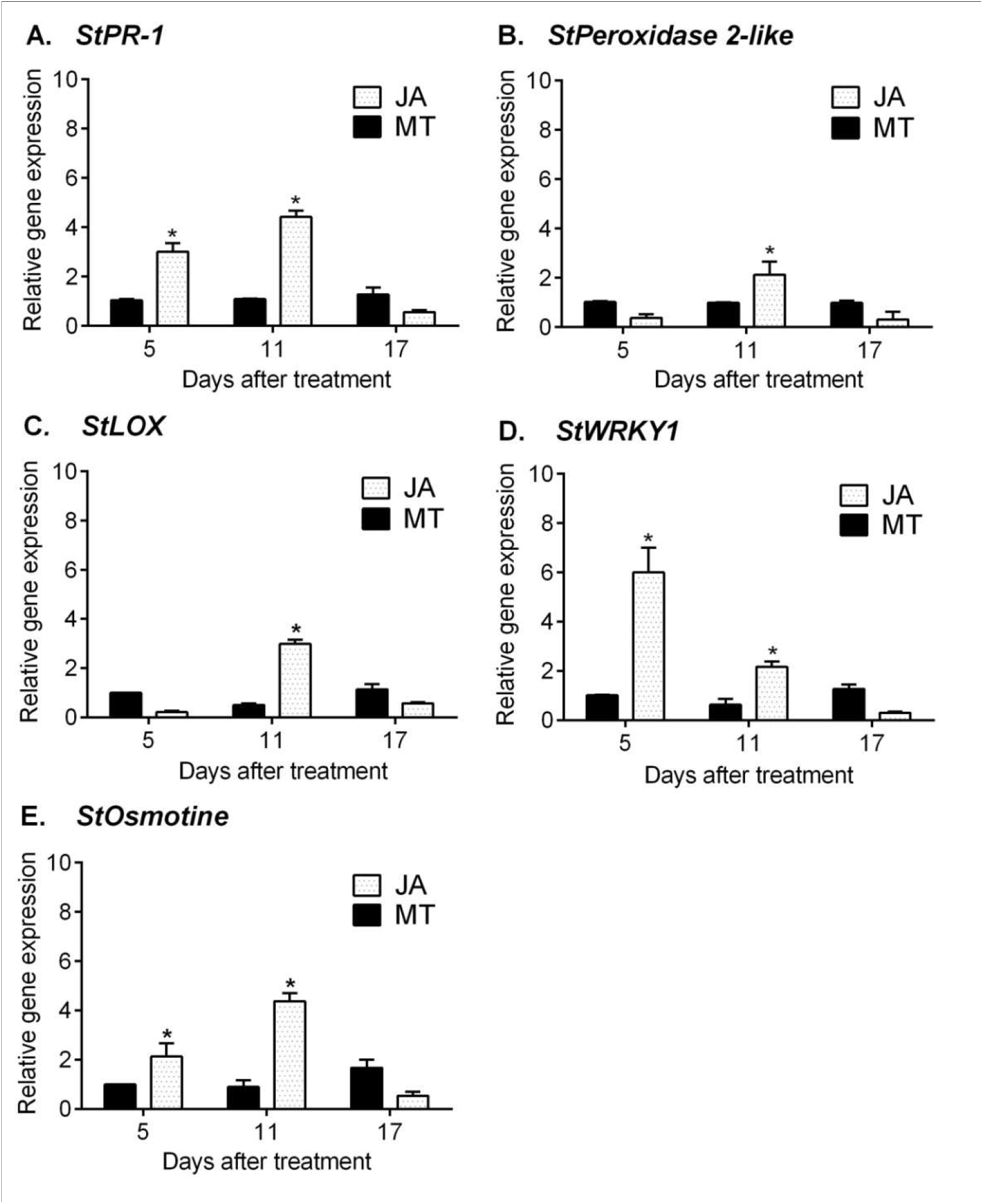
Relative transcriptional expression of defense response-related genes in potato treated with JA or MT and inoculated with *P. infestans* 5, 11 and 17 days after treatment (DAT). **(A)** *StPR1*, **(B)** *StPeroxidase 2-like*, **(C)** *StLox*, **(D)** *StWRKY1*, **(E)** *StOsmotine*. Samples were taken 48 hours post-inoculation. Expression was normalized with housekeeping *StEF1*α and relative expression levels were calculated using MT-5DAT treatment as a reference. The bars represent the mean ± SE. Values represent means from at least three independent experiments. Significant differences in relative expression levels using the t-Student, *: Significant at level 0.05.

In addition, the expression of other defense-response genes was also analyzed. Genes encoding *StPeroxidase* (Figure 8B) and *StLOX* (Figure 8C) showed significantly higher relative expression (P <0.05) only in JA-treated plants inoculated 11DAT, with expression levels being two (p=0.034*) and three (p=0.015*) times higher, respectively, compared to MT. Genes coding for *StWRKY* (Figure 8D) and *StOsmotine* (Figure 8E) showed significantly higher relative expression (P <0.05) in JA-treated plants inoculated 5DAT and 11DAT, with expression levels six (p=0.013*) and two (p=0.028*) times higher and two (p=0.032*) and 4.5 (p=0.011*) times higher, respectively, compared to MT. At 17 days, no significant differences were found between treated plants with JA or MT. These results demonstrate that the protective effect of JA against *P. infestans* is associated with increased expression of salycilic acid-dependent and independent defense-related genes.

## 4. Discussion

In this study, we show that JA transiently modifies the physiological status of potato and generates a priming state as *StMYC2* expression diminishes. By means of the gene expression, physiological and microscopy analysis, we observed a protective effect 11 DAT in JA-treated plants against *P. infestans* probably achieved by an increased transcription of defense genes, decreasing the abundance of developed structures and delaying the progression of the disease.

The JA concentration used (150 µg/mL ∼0.72 µM) generated a transitory alteration of the physiological state of the plant, prior to entering the primed state. The initial effect of the hormone induced *StMYC2* expression at least during the first five days (Figure 6), in parallel with partial stomatal closure, which lasted 10-12 days (Figure 1B). Jasmonates have been previously reported to regulate stomatal closure [56]; the observed stomatal conductance response is probably a direct effect produced by JA and consequently can be considered as a non-invasive indicator of JA-mediated signaling in plants. Regarding Fv/Fm values, close to 0.835 in our experiment, no significant differences were observed between treatments (Figure 1A). This suggests that JA does not alter the light absorbing efficiency PSII photochemistry. This parameter has been used as an indicator of physiological stress, as decreases might reflect a lower efficiency of PSII [39, 57]. Values close to 0.835 are considered ideal for a large number of species [58, 59].

Five days after JA treatment, an strong transcription of *StMYC2* was observed with and without *P. infestans* inoculation (Figures 6 and 7A), and a slight increase in transcription of only some defense genes associated with response mediated by SA (Figure 8). This observation demonstrates that *StMYC2* expression is not responding to *P. infestans* inoculation. Previous studies have shown that transcription of *MYC2* hinders the expression of diverses defense genes, thus impeding plant resistance to various pathogens [46,47,55,60]. The increase of *StMYC2* expression at 5DAT possibly interfered with a correct activation of the defense response against *P. infestans*. But interestingly, an improved transcription only of *StPR1* and *StOsmotine* was evidenced at 5 DAT, both actively involved in HR [49,61,62,63], and of the transcription factor *StWRKY1* involved in the regulation of genes related to the strengthening of the cell wall [45]. Probably, the activation of these genes is associated with the reduction of necrotic lesions that became evident after 72 hours at 5 DAT (Figure 5A). Taken together, results show that the active presence of *StMYC2* at 5DAT dampens the defense response against *P. infestans*.

Macroscopic, physiological and microscopic analyses showed the priming effect of JA, 11 days after treatment, against *P. infestans*. We observed a delay in the progression of the disease (Figure 2) and reduced physiological stress caused by *P. infestans* compared to the other treatments (Figures 3A, B). These observations were corroborated with a decrease of pathogen structures (Figure 4) and a significant reduction in the number of necrotic lesions at 48 and 72 hpi (Figure 5B). Differences in abundance of structures and the delay in the initiation of the necrotrophic phase in *P. infestans* have been previously demonstrated in tomato and potato cultivars with partially compatible and strong compatible interactions, implying that there is a modulation in the type of interaction when there is an enhanced defense response in plants [44,64,65].

At the expression level, a stronger activation of *StPR1, StOsmotine* and *StWRKY1* was observed in response to *P. infestans* 11DAT, compared to the expression observed at 5DAT, along with gene activation of *StPeroxidase-2Like* and *StLipoxigenase* (Figure 8). These genes are involved in lignin accumulation, ROS control and suberization, respectively [66,67]. This incremented expression might be associated to the reduction in *StMYC2* accumulation at 11DAT (Figure 6), as MYC2 expression has been related with hampering of SA-dependent defense [68]. These results suggest that JA-dependent priming is active at 11DAT by preparing and potentiating the transcription of genes related to HR, ROS and cell wall reinforcement against *P. infestans*.

At 17 days after JA treatment, no protective effect was observed mediated by the application of JA. No significant differences were detected in defense gene transcription (Figure 8), Fv/Fm values, providing evidence of the stress generated by the pathogenic progression of *P. infestans* (Figure 3A), and number of necrotic lesions (Figure 5C), which would suggest that the priming effect generated by JA has ended.

Cohen et al. [31] previously reported a protective effect of JA in potato and tomato in response to *P. infestans* from 5DAT, and its termination 11DAT. They also showed that concentration has an effect on the percentage of protection over time. In our study we observed at 5DAT that *StMYC2-*mediated signaling probably interferes with the protective effect. This observation may be a consequence of a higher sensitivity to JA of the genotype tested (Criolla Colombia, a diploid cultivar). Our results suggest that the generation and duration of the JA-priming effect is dependent not only on the concentration, but also on the the genotype of the plant as previously reported with BABA [69]. Subsequent investigations could evaluate the optimal JA concentration to generate a faster and longer-lasting priming effect at a local and systemic level.

Based on our results, we conclude that the priming effect generated by the application of JA acts within a specific time window (Figure 9). Five days after treatment the initial plant reaction to JA is evidenced by stomatal closure and activation of *StMYC2*, which possibily obstructs the priming effect. 11DAT, a significant priming effect is observed, associated with increased transcriptional induction of genes involved in different response pathways, which especially impact *P. infestans* change to necrotrophy. After 17 days the priming effect of JA ends, the plant again shows susceptibility to *P. infestans* and no differences were observed in any of the variables evaluated between treatments.

**Figure 9.**
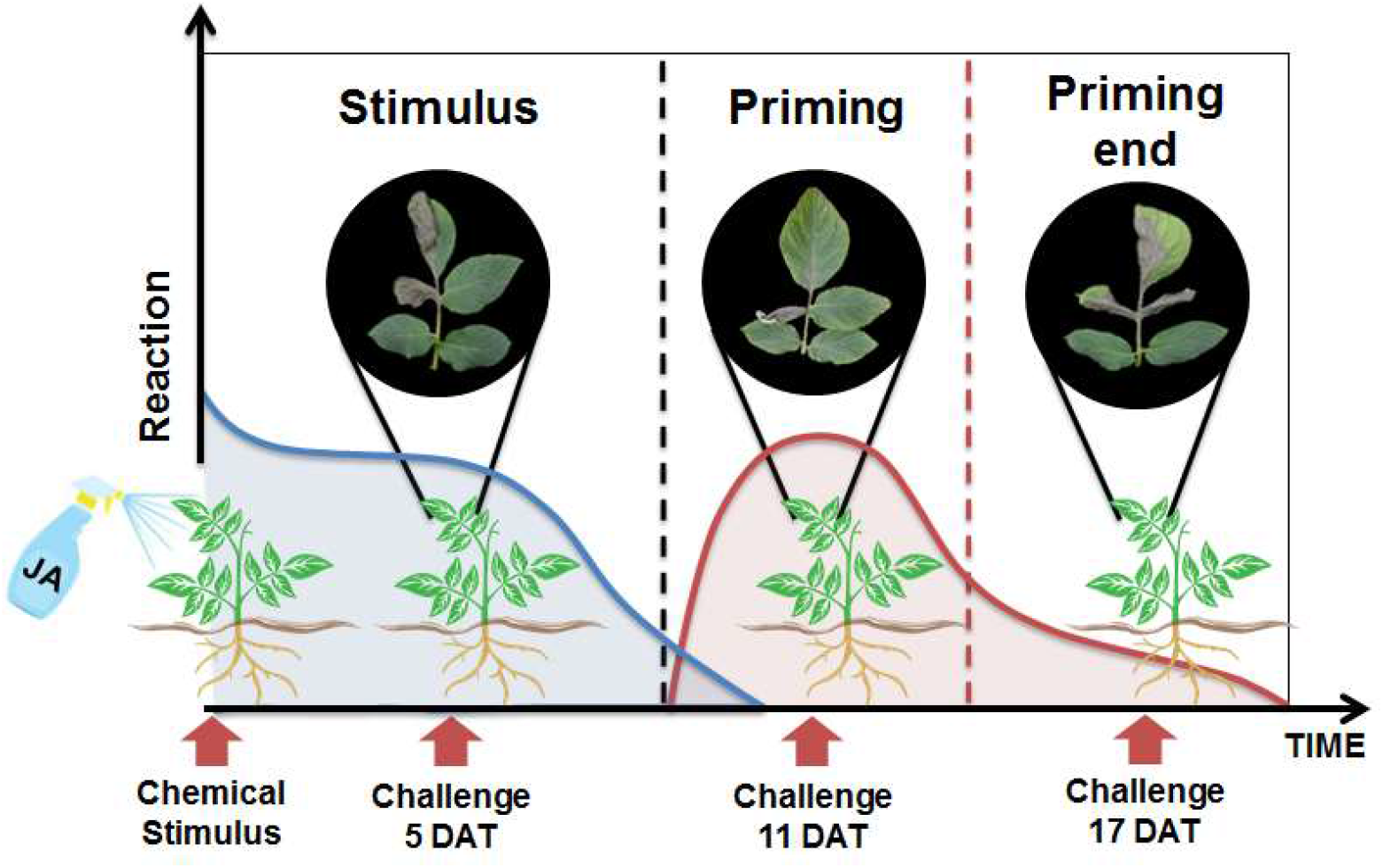
Priming effect JA in potato against *P. infestans* acts within a specific time window. The blue line represents the transcriptional activation of *StMYC2* and the change in stomatal conductance generated by the JA treatment. The red line represents the level of reaction and the duration of the priming effect produced by the stimulus.

The molecular basis of priming has been associated to histone modification, accumulation/protection of mRNAs, accumulation of protein kinases, DNA methylation and activation of hormone biosynthesis [20,21,70]. Previous studies have shown that exogenous application of BABA induces transgenerational resistance in potato to *P. infestans* by priming expression of defense genes sensitive to SA, altering DNA methylation and histone modification patterns [71,72]. Likewise, exogenous application of JA induces tolerance to dehydration stress in *Arabidopsis* by priming the transcription of ABA-dependent genes. Histone modification also seems to be the molecular mechanism involved in the priming effect [73]. These mechanisms could be involved in the protective effect generated by JA in potato; experiments associated with transcriptome analysis and chromatin configuration in potato could determine the molecular basis behind the priming observed at 11 days.

## Nomenclature

JA= Jasmonic acid

hpi= Hours post inoculation

DAT = Days after treatment

## Acknowledgements

The authors thank Professor Luis Fernando Cadavid Gutiérrez and the laboratory assistant Blanca Elvira Schroeder López from the Instituto de Genética at Universidad Nacional de Colombia for the training and support to carry out the gene expression experiments. The authors also wish to acknowledge Chary Esteban Quinche González and Ivon Arcila for the collaboration with the microscopy analysis and plant inoculation, respectively, and Professors Joaquin Ramirez and Elena Brochero from the Facultad de Ciencias Agrarias for all the revisions and useful suggestions.

## References

[1] Food and Agriculture Organization of the United Nations. (2018). World food and agriculture: Statistical pocketbook 2018.

[2] Fry, W. E., Birch, P. R. J., Judelson, H. S., Grünwald, N. J., Danies, G., Everts, K. L., et al. (2015). Five Reasons to Consider Phytophthora infestans a Reemerging Pathogen. Phytopathology. 105, 966–981. doi: 10.1094/PHYTO-01-15-0005-FI

[3] Kamoun, S., Furzer, O., Jones, J. D. G., Judelson, H. S., Ali, G. S., Dalio, R. J. et al. (2015). The Top 10 oomycete pathogens in molecular plant pathology. Molecular Plant Pathology. 16, 413–434. doi.org/10.1111/mpp.12190

[4] Hassan, M. A., and Abo-Elyousr, K. A. (2013). Activation of tomato plant defence responses against bacterial wilt caused by Ralstonia solanacearum using DL-3-aminobutyric acid (BABA). European Journal of Plant Pathology. 136, 145–157. doi:10.1007/s10658-012-0149-4

[5] Haverkort, A. J., Boonekamp, P. M., Hutten, R., Jacobsen, E., Lotz, L. a. P., Kessel, G. J. et al. (2016). Durable Late Blight Resistance in Potato Through Dynamic Varieties Obtained by Cisgenesis: Scientific and Societal Advances in the DuRPh Project. Potato Research. 35–66. doi.org/10.1007/s11540-015-9312-6

[6] Kamoun, S., and Smart, C. D. (2005). Late Blight of Potato and Tomato in the Genomics Era. Plant Disease. 89, 692–699. doi; 10.1094/PD-89-0692

[7] Agrios, G. N. (2005). Plant pathology (5th ed). Elsevier Academic Press.

[8] Delalieux, S., van Aardt, J., Keulemans, W., Schrevens, E., and Coppin, P. (2007). Detection of biotic stress (Venturia inaequalis) in apple trees using hyperspectral data: Non-parametric statistical approaches and physiological implications. European Journal of Agronomy. 27, 130–143. doi: 10.1016/j.eja.2007.02.005

[9] Rubio, J., López, C., and Melgarejo, L. M. (2017). Physiological behavior of cassava plants (Manihot esculenta Crantz) in response to infection by Xanthomonas axonopodis pv. Manihotis under greenhouse conditions. Physiological and Molecular Plant Pathology. 100. doi: 10.1016/j.pmpp.2017.09.004

[10] Selvaraj, K., and Fofana, B. (2012). An Overview of Plant Photosynthesis Modulation by Pathogen Attacks. Advances in Photosynthesis – Fundamental Aspects. 22, 466–484. doi: 10.5772/27124

[11] Robert-Seilaniantz, A., Grant, M., and Jones, J. D. G. (2011). Hormone crosstalk in plant disease and defense: More than just jasmonate-salicylate antagonism. Annual Review of Phytopathology. 49, 317–343. doi: 10.1146/annurev-phyto-073009-114447

[12] Glazebrook, J. (2005). Contrasting mechanisms of defense against biotrophic and necrotrophic pathogens. Annual Review of Phytopathology. 43, 205–227. doi: 10.1146/annurev.phyto.43.040204.135923

[13] Halim, V. A., Vess, A., Scheel, D., and Rosahl, S. (2006). The role of salicylic acid and jasmonic acid in pathogen defence. Plant Biology (Stuttgart, Germany). 8, 307–313. doi: 10.1055/s-2006-924025

[14] Pieterse, C. M. J., Van der Does, D., Zamioudis, C., Leon-Reyes, A., and Van Wees, S. C. M. (2012). Hormonal modulation of plant immunity. Annual Review of Cell and Developmental Biology. 28, 489–521. doi: 10.1146/annurev-cellbio-092910-154055

[15] Jia, X., Zeng, H., Wang, W., Zhang, F., and Yin, H. (2018). Chitosan Oligosaccharide Induces Resistance to Pseudomonas syringae pv. Tomato DC3000 in Arabidopsis thaliana by Activating Both Salicylic Acid– and Jasmonic Acid–Mediated Pathways. Molecular Plant-Microbe Interactions®. 31, 1271–1279. doi: 10.1094/MPMI-03-18-0071-R

[16] Thaler, J. S., Owen, B., and Higgins, V. J. (2004). The role of the jasmonate response in plant susceptibility to diverse pathogens with a range of lifestyles. Plant Physiology. 135, 530–538. doi: 10.1104/pp.104.041566

[17] Castaño Monsalve, J., Ramírez Gil, J.G. Patiño Hoyos, L. F., and Morales Osorio, J. G. (2015). Alternativa para el manejo de Phytophthora infestans (Mont.) de Bary en Solanum betaceum Cav. Mediante inductores de resistencia. Revista de Protección Vegetal. 30, 204–212.

[18] Król, P., Igielski, R., Pollmann, S., and Kępczyńska, E. (2015). Priming of seeds with methyl jasmonate induced resistance to hemi-biotroph Fusarium oxysporum f.sp. Lycopersici in tomato via 12-oxo-phytodienoic acid, salicylic acid, and flavonol accumulation. Journal of Plant Physiology. 179, 122–132. doi: 10.1016/j.jplph.2015.01.018

[19] Pajerowska-Mukhtar, K., Stich, B., Achenbach, U., Ballvora, A., Lübeck, J., Strahwald, J., et al. (2009). Single Nucleotide Polymorphisms in the Allene Oxide Synthase 2 Gene Are Associated With Field Resistance to Late Blight in Populations of Tetraploid Potato Cultivars. Genetics. 181, 1115–1127. doi: 10.1534/genetics.108.094268

[20] Balmer, A., Pastor, V., Gamir, J., Flors, V., and Mauch-Mani, B. (2015). The “prime-ome”: Towards a holistic approach to priming. Trends in Plant Science. 20, 443–452. doi: 10.1016/j.tplants.2015.04.002

[21] Crisp, P. A., Ganguly, D., Eichten, S. R., Borevitz, J. O., and Pogson, B. J. (2016). Reconsidering plant memory: Intersections between stress recovery, RNA turnover, and epigenetics. Science Advances. 2, 1–14. doi: 10.1126/sciadv.1501340

[22] Zhou, M., and Memelink, J. (2016). Jasmonate-responsive transcription factors regulating plant secondary metabolism. Biotechnology Advances. 34, 441–449. doi: 10.1016/j.biotechadv.2016.02.004

[23] Martinez-Medina, A., Flors, V., Heil, M., Mauch-Mani, B., Pieterse, C. M. J., Pozo, M. J., et al. (2016). Recognizing Plant Defense Priming. Trends in Plant Science. 21, 818– 822. doi: 10.1016/j.tplants.2016.07.009

[24] Gamir, J., Sánchez-Bel, P., and Flors, V. (2014). Molecular and physiological stages of priming: how plants prepare for environmental challenges. Plant Cell Reports. 33, 1935–1949. doi:10.1007/s00299-014-1665-9

[25] Pastor, V., Balmer, A., Gamir, J., Flors, V., and Mauch-Mani, B. (2014). Preparing to fight back: generation and storage of priming compounds. Frontiers in Plant Science. 5, 295. doi:10.3389/fpls.2014.00295

[26] Aranega-Bou P, delaO Leyva M, Finiti I, Garcia-Agistin P, Gonzalez-Bosch C. (2014). Priming of plant resistance by natural compounds. Hexanoic acid as a model. Frontiers in Plant Science. 5:488. doi: 10.3389/fpls.2014.00488

[27] Ávila, A. C., Ochoa, J., Proaño, K., and Martínez, M. C. (2019). Jasmonic acid and nitric oxide protects naranjilla (Solanum quitoense) against infection by Fusarium oxysporum f. Sp. Quitoense by eliciting plant defense responses. Physiological and Molecular Plant Pathology. 106, 129–136. doi: 10.1016/j.pmpp.2019.01.002

[28] Ueeda, M., Kubota, M., and Nishi, K. (2005). Contribution of jasmonic acid to resistance against Phytophthora blight in Capsicum annuum cv. SCM334. Physiological and Molecular Plant Pathology. 67, 149–154. doi: 10.1016/j.pmpp.2005.12.002

[29] Floryszak-Wieczorek, J., Magdalena, A.-J. M., & Abramowski, D. (2015). BABA-primed defense responses to Phytophthora infestans in the next vegetative progeny of potato. Frontiers in Plant Science. 6, 844. doi:10.3389/fpls.2015.00844.

[30] Val, F., Desender, S., Bernard, K., Potin, P., Hamelin, G., and Andrivon, D. (2008). A culture filtrate of Phytophthora infestans primes defense reaction in potato cell suspensions. Phytopathology, 98(6), 653–658. doi: 10.1094/PHYTO-98-6-0653

[31] Cohen, Y., Gisi, U., and Niderman, T. (1993). Local and Systemic Protection Against Phytophthora infestans Induced in Potato and Tomato Plants by Jasmonic Acid and Jasmonic Methyl Ester. Phytopathology. 83. doi: 10.1094/Phyto-83-1054

[32] Walters, D and Heil, M (2007) Costs and trade-offs associated with induced resistance. Physiological and Molecular Plant Pathology. 71, 3–17. doi: 10.1016/j.pmpp.2007.09.008

[33] Yogendra, K. N., Kushalappa, A. C., Sarmiento, F., Rodriguez, E., and Mosquera, T. (2015b). Metabolomics deciphers quantitative resistance mechanisms in diploid potato clones against late blight. Functional Plant Biology. 42, 284–298. doi: 10.1071/FP14177

[34] Meier, U. (1997). Growth stages of mono-and dicotyledonous plants. Blackwell Wissenschafts-Verlag.

[35] Caten, C. E., and Jinks, J. L. (1968). Spontaneous variability of single isolates of Phytophthora infestans. I. Cultural variation. Canadian Journal of Botany. 46, 329–348. doi: 10.1139/b68-055

[36] García, H. G., Marín, M., Jaramillo, S., and Cotes, J. M. (2008). Sensitivity to four systemic fungicides of Colombian isolates of Phytophthora infestans. Agronomía Colombiana. 26, 47–57.

[37] Yogendra KN, Pushpa D, Mosa KA, Kushalappa AC, Murphy A, Mosquera T. (2014). Quantitative resistance in potato leaves to late blight associated with induced hydroxycinnamic acid amides. Functional & Integrative Genomics. 5, 285–298. doi: 10.1007/s10142-013-0358-8

[38] Gyetvai G., Sønderkær M., Göbel U., Basekow R., Ballvora A., Imhoff M., et al.. (2012). The transcriptome of compatible and incompatible interactions of potato (Solanum atuberosum) with Phytophthora infestans revealed by DeepSAGE analysis. PLoS ONE. 7, e31526. doi: 10.1371/journal.pone.0031526

[39] Murchie, E.H., and Lawson, T. (2013). Chlorophyll fluorescence analysis: a guide to good practice and understanding some new applications. Journal of experimental botany. 64, 3983–3998. doi: 10.1093/jxb/ert208

[40] Campbell, C. L., and Madden, L. V. (1990). Introduction to plant disease epidemiology. Wiley, New York. 1990.p. 73.

[41] Bonierbale, M., Haan, S., Forbes, A., and Bastos, C. (2010). Procedimientos para pruebas de evaluacion estandar de clones avanzados de papa: Guia para cooperadores internacionales. Lima (Peru). Centro Internacional de la Papa (CIP). 151p.

[42] Chung, C.-L., Longfellow, J. M., Walsh, E. K., Kerdieh, Z., Van Esbroeck, G., Balint-Kurti, P., et al. (2010). Resistance loci affecting distinct stages of fungal pathogenesis: Use of introgression lines for QTL mapping and characterization in the maize--Setosphaeria turcica pathosystem. BMC Plant Biology. 10, 103. doi: 10.1186/1471-2229-10-103

[43] Knox-Davies, P. S. (1974). Penetration of maize leaves by Helminthosporium turcicum. Phytopathology. 64, 1468–1470.

[44] Zuluaga, A. P., Vega-Arreguín, J. C., Fei, Z., Matas, A. J., Patev, S., Fry, W. E., et al. (2016). Analysis of the tomato leaf transcriptome during successive hemibiotrophic stages of a compatible interaction with the oomycete pathogen Phytophthora infestans. Molecular Plant Pathology. 17, 42–54. doi: 10.1111/mpp.12260

[45] Yogendra, K. N., Kumar, A., Sarkar, K., Li, Y., Pushpa, D., Mosa, K. A., Duggavathi, R., et al. (2015a). Transcription factor StWRKY1 regulates phenylpropanoid metabolites conferring late blight resistance in potato. Journal of Experimental Botany. 66, 7377–7389. doi: 10.1093/jxb/erv434

[46] Boter, M., Ruíz-Rivero, O., Abdeen, A., and Prat, S. (2004). Conserved MYC transcription factors play a key role in jasmonate signaling both in tomato and Arabidopsis. Genes & Development. 18, 1577–1591. doi: 10.1101/gad.297704

[47] Du, M., Zhao, J., Tzeng, D. T. W., Liu, Y., Deng, L., Yang, T., et al. (2017). MYC2 Orchestrates a Hierarchical Transcriptional Cascade That Regulates Jasmonate-Mediated Plant Immunity in Tomato. The Plant Cell. 29, 1883–1906. doi: 10.1105/tpc.16.00953

[48] Gallou, A., Cranenbrouck, S., Declerck, S., Gallou, A., Cranenbrouck, S., and Declerck, S (2009). Trichoderma harzianum elicits defence response genes in roots of potato plantlets challenged by Rhizoctonia solani. European Journal of Plant Pathology. 124, 219–230. doi:org/10.1007/s10658-008-9407-x

[49] Orłowska, E., Fiil, A., Kirk, H.-G., Llorente, B., and Cvitanich, C. (2012). Differential gene induction in resistant and susceptible potato cultivars at early stages of infection by Phytophthora infestans. Plant Cell Reports. 31, 187–203. doi: 10.1007/s00299-011-1155-2

[50] Shoji, T., and Hashimoto, T. (2011). Tobacco MYC2 regulates jasmonate-inducible nicotine biosynthesis genes directly and by way of the NIC2-locus ERF genes. Plant & Cell Physiology. 52, 1117–1130. doi: 10.1093/pcp/pcr063

[51] Hossain, M. A., Munemasa, S., Uraji, M., Nakamura, Y., Mori, I. C., and Murata, Y. (2011). Involvement of endogenous abscisic acid in methyl jasmonate-induced stomatal closure in Arabidopsis. Plant Physiology. 156, 430–438. doi: 10.1104/pp.111.172254

[52] Nicot, N., Hausman, J.-F., Hoffmann, L., and Evers, D. (2005). Housekeeping gene selection for real-time RT-PCR normalization in potato during biotic and abiotic stress. Journal of Experimental Botany. 56, 2907–2914. doi: 10.1093/jxb/eri285

[53] Muller, P. Y., Janovjak, H., Miserez, A. R., and Dobbie, Z. (2002). Processing of gene expression data generated by quantitative real-time RT-PCR. BioTechniques. 32, 1372–1374, 1376, 1378–1379.

[54] Simon, P. (2003). Q-Gene: Processing quantitative real-time RT-PCR data. Bioinformatics (Oxford, England). 19, 1439–1440. doi: 10.1093/bioinformatics/btg157

[55] Dombrecht, B., Xue, G. P., Sprague, S. J., Kirkegaard, J. A., Ross, J. J., Reid, J. B., et al. (2007). MYC2 differentially modulates diverse jasmonate-dependent functions in Arabidopsis. The Plant Cell. 19, 2225–2245. doi: 10.1105/tpc.106.048017

[56] Behiry, S; Ashmawy, N; Abdelkhalek, A; Khaled, A and Elsayed H. (2017). “Compatible and incompatible type interactions related to defense genes in Potato elucidation by Pectobacterium carotovorum”. Journal of Plant Diseases and Protection -New Series-. 125. doi:10.1007/s41348-017-0125-5

[57] Baker, N. R. (2008). Chlorophyll fluorescence: A probe of photosynthesis in vivo. Annual Review of Plant Biology. 59, 89–113. doi: 10.1146/annurev.arplant.59.032607.092759

[58] Björkman, O., and Demmig, B. (1987). Photon yield of O2 evolution and chlorophyll fluorescence characteristics at 77 K among vascular plants of diverse origins. Planta. 170, 489–504. doi: 10.1007/BF00402983

[59] Roháček, K., Soukupová, J., and Bartak, M. (2008). Chlorophyll fluorescence: A wonderful tool to study plant physiology and plant stress. Plant Cell Compartments - Selected Topics. 41, 104.

[60] Lorenzo, O., Chico, J. M., Sánchez-Serrano, J. J., and Solano, R. (2004). JASMONATE-INSENSITIVE1 encodes a MYC transcription factor essential to discriminate between different jasmonate-regulated defense responses in Arabidopsis. The Plant Cell. 16, 1938–1950. doi: 10.1105/tpc.022319

[61] Anil Kumar, S., Hima Kumari, P., Shravan Kumar, G., Mohanalatha, C., and Kavi Kishor, P. B. (2015). Osmotin: A plant sentinel and a possible agonist of mammalian adiponectin. Frontiers in Plant Science. 6. doi: 10.3389/fpls.2015.00163

[62] Hao, D., Yang, J., Long, W., Yi, J., VanderZaag, P., and Li, C. (2018). Multiple R genes and phenolic compounds synthesis involved in the durable resistance to Phytophthora infestans in potato cv. Cooperation 88. Agri Gene. 8, 28–36. doi: 10.1016/j.aggene.2018.04.001

[63] Viktorova, J., Krasny, L., Kamlar, M., Sura-de Jong, M., Mackova, M., and Macek, T. (2012). Osmotin, a Pathogenesis-Related Protein. Current Protein & Peptide Science. 13. doi: 10.2174/138920312804142129

[64] Cai, G., Restrepo, S., Myers, K., Zuluaga, P., Danies, G., Smart, C., et al. (2012). Gene profiling in partially resistant and susceptible near-Isogenic tomatoes in response to late blight in the field. Molecular Plant Pathology. 14. doi: 10.1111/j.1364-3703.2012.00841.x

[65] Vega-Sánchez, M. E., Erselius, L. J., Rodriguez, A. M., Bastidas, O., Hohl, H. R., Ojiambo, P. S., et al. (2000). Host adaptation to potato and tomato within the US-1 clonal lineage of Phytophthora infestans in Uganda and Kenya. Plant Pathology. 49, 531–539. doi: 10.1046/j.1365-3059.2000.00487.x

[66] Sorokan, A. V., Burhanova, G. F., and Maksimov, I. V. (2018). Anionic peroxidase-mediated oxidative burst requirement for jasmonic acid-dependent Solanum tuberosum defence against Phytophthora infestans. Plant Pathology. 67, 349–357. doi: 10.1111/ppa.12743

[67] Tejeda-Sartorius, M., and Martínez-Gallardo, N. (2007). Jasmonic Acid Accelerates the Expression of a Pathogen-Specific Lipoxygenase (POTLX-3) and Delays Foliar Late Blight Development in Potato (Solanum tuberosum L.). Revista Mexicana de Fitopatologia. 25, 18–25.

[68] Zheng, X. Y., Spivey, N. W., Zeng, W., Liu, P. P., Fu, Z. Q., Klessig, D. F. …, & Dong, X. (2012). Coronatine promotes Pseudomonas syringae virulence in plants by activating a signaling cascade that inhibits salicylic acid accumulation. Cell host & microbe, 11(6), 587–596.

[69] Bengtsson, T., Holefors, A., Witzell, J., Andreasson, E., and Liljeroth, E. (2013). Activation of defence responses to Phytophthora infestansin potato by BABA. Plant Pathology, 63(1), 193–202. doi:10.1111/ppa.12069

[70] Conrath, U., Beckers, G. J. M., Langenbach, C. J. G., and Jaskiewicz, M. R. (2015). Priming for enhanced defense. Annual Review of Phytopathology. 53, 97–119. doi: 10.1146/annurev-phyto-080614-120132

[71] Meller, B., Kuźnicki, D., Arasimowicz-Jelonek, M., Deckert, J., and Floryszak-Wieczorek, J. (2018). BABA-Primed Histone Modifications in Potato for Intergenerational Resistance to Phytophthora infestans. Frontiers in Plant Science. 9, 1228. doi: 10.3389/fpls.2018.01228

[72] Kuźnicki, D., Meller, B., Arasimowicz-Jelonek, M., Braszewska-Zalewska, A., Drozda, A., and Floryszak-Wieczorek, J. (2019). BABA-Induced DNA Methylome Adjustment to Intergenerational Defense Priming in Potato to Phytophthora infestans. Frontiers in Plant Science. 10. doi: 10.3389/fpls.2019.00650

[73] Liu, N., Staswick, P. E., and Avramova, Z. (2016). Memory responses of jasmonic acid-associated Arabidopsis genes to a repeated dehydration stress. Plant, Cell & Environment. 39, 2515–2529. doi: 10.1111/pce.12806

